# Krisp: A python package for designing CRISPR and amplification-based diagnostic assays from whole genome data

**DOI:** 10.1101/2023.11.16.567433

**Authors:** Zachary S. L. Foster, Andrew S. Tupper, Caroline M. Press, Niklaus J. Grünwald

## Abstract

Recent pandemics such as COVID-19 have highlighted the importance of rapidly developing diagnostics to detect and monitor evolving pathogens. CRISPR-Cas technology, combined with isothermal DNA amplification methods, has recently been used to develop diagnostic assays for sequence-specific recognition of DNA or RNA. These assays have similar sensitivity to the gold standard qPCR but can be deployed as easy to use and inexpensive test strips. However, the discovery of diagnostic regions of a genome flanked by conserved regions where primers can be designed requires extensive bioinformatic analyses of genome sequences. We developed the python package `krisp` to find primers and diagnostic sequences that differentiate groups of samples from each other at any taxonomic scale, using either unaligned genome sequences or a variant call format (VCF) file as input. `Krisp` has been optimized to handle large datasets by using efficient algorithms that run in near linear time, use minimal RAM, and leverage parallel processing when available. The validity of `krisp` results has been demonstrated in the laboratory with the successful design of SHERLOCK assays to distinguish the sudden oak death pathogen *Phytophthora ramorum* from closely related *Phytophthora* species. `Krisp` is released open source under a permissive license with all the documentation needed to quickly design CRISPR-Cas diagnostic assays.

**Author summary:** Pathogens continue to emerge at accelerated rates affecting animals, plants, and ecosystems. Rapid development of novel diagnostic tools is needed to monitor novel pathogen variants or groups. We developed the computational tool `krisp` to identify genetic regions suitable for development of CRISPR diagnostics and traditional amplification-based diagnostics such as PCR. `Krisp` scans whole genome sequence data for target and non-target groups to identify diagnostic regions based on DNA or RNA sequences. This computational tool has been validated using genome data for the sudden oak death pathogen *Phytophthora ramorum*. `Krisp` is released open source under a permissive license with all the documentation needed to quickly design CRISPR-Cas diagnostic assays and other amplification-based assays.

## Introduction

Invasive organisms continue to emerge at accelerated rates worldwide, likely due to increased global trade, movement of live plants, a growing human population, and stresses imposed by the Anthropocene on hosts (Fisher et al., 2012; Hulme, 2009; Liebhold et al., 2012; Meyerson & Mooney, 2007; Waters et al., 2016). Most pathogens, whether bacteria, fungi, oomycetes, viruses, nematodes, or other protists, are hard to detect without appropriate diagnostics. The biosurveillance of the future will require affordable, robust, accurate, and field deployable diagnostic methods that can be developed quickly. The capacity to develop novel diagnostics in response to an emergent pathogen in weeks, rather than months or years, would be transformative. Whole genome sequences (WGS) of new variants combined with computational tools to identify diagnostic loci provides a new means of accelerating developing diagnostics in response to emerging or reemerging invasive pathogens and pests.

Clustered regularly interspaced short palindromic repeats (CRISPR) are DNA sequences found in prokaryotes to detect and counter viral infections (Barrangou, 2015). These loci consist of a series of conserved repeats alternating with equal length variable sequences referred to as spacers. The spacers are sequences copied from viruses that the microbe or its progenitors have encountered in the past. CRISPR-associated proteins (Cas) use RNA transcribed from the spacer sequences as guides to recognize and, depending on the CAS enzyme, degrade specific strands of DNA or RNA. For example, the Cas9 protein introduces double-stranded breaks near DNA sequences complementary to the bound guide RNA (Jinek et al., 2012). In addition to sequence-specific cleavage, the Cas12, Cas13, and Cas14 families of proteins cause cleavage of off-target nucleic acid polymers after binding to and cleaving a target sequence, a phenomena referred to as collateral cleavage (Abudayyeh et al., 2016; Chen et al., 2018; East-Seletsky et al., 2016; Harrington et al., 2018). For Cas13 orthologs, both sequence-specific cleavage and collateral cleavage target RNA (Gootenberg et al., 2018), whereas Cas12 and the smaller Cas14 variant target ssDNA or dsDNA and cleave bystander ssDNA (Chen et al., 2018; Harrington et al., 2018). Cas enzymes also differ in regard to which 2-6bp sequence motifs are most effected by collateral cleavage (Gootenberg et al., 2018). As part of the bacterial immune system, collateral cleavage is thought to limit pathogen propagation via programmed cell death or dormancy induction (Abudayyeh et al., 2016, p. 2), thereby sacrificing the individual to benefit the population as a whole.

The discovery and understanding of the CRIPSR-Cas immune system has led to many practical advances in molecular biology, including CRISPR-based diagnostic assays for detecting specific RNA or DNA sequences (CRISPR-dx). Many techniques rely on using the collateral cleavage activity of the Cas12, Cas13, and Cas14 enzymes to degrade ssDNA or RNA reporters as a measurable signal of sequence recognition. The degradation of reporter molecules can be detected by a variety of ways, including measuring fluorescence, imaging fluorescent bands on a gel, or observing bands on a lateral flow device (Kellner et al., 2019). In the case of solution-based fluorometric detection, short off-target RNA or ssDNA polymers, each with a fluorescent probe on one end and a quencher on the other, are cut by the collateral cleavage of Cas proteins upon sequence recognition, resulting in fluorescence (East-Seletsky et al., 2016). This general technique has been used to create highly specific CRISPR-based diagnostic assays. SHERLOCK (Kellner et al., 2019) and DETECTR (Chen et al., 2018) achieve high sensitivity using amplification of target DNA by recombinase polymerase amplification (RPA), an isothermal alternative to PCR (Daher et al., 2016). HOLMES (Li et al., 2019) uses loop-mediated isothermal amplification (LAMP), another isothermal amplification technique (Notomi et al., 2000). CONAN achieves high sensitivity without preamplification by using a sophisticated DNA/RNA hybrid reporter that, when degraded, releases a guide RNA that matches a fragment of DNA added to the reaction, resulting in exponential activation of the Cas12a enzyme (Shi et al., 2021). These assays have been shown to be both highly specific, allowing single nucleotide discrimination, and very sensitive, with detection thresholds in the attomolar range (Chen et al., 2018; Gootenberg et al., 2018). Sensitivity can be further increased by scaling up preamplification or adding multiple guide RNAs targeting different parts of the same target sequence into the same reaction (Fozouni et al., 2021). These methods can be adapted to single-tube isothermal reactions or lateral flow strips, allowing for on-site detection of pathogens with minimal specialized instruments or training for as little as $0.60 per assay (Kellner et al., 2019). Other CRISPR-dx techniques involve engineered tracrRNAs to make the target RNA act as a guide RNA (LEOPARD) (Jiao et al., 2021), observing liquid-liquid phase separation caused by collateral cleavage (Spoelstra et al., 2021), engineered hydrogels that change their material properties in response to a target (English et al., 2019), measuring the byproducts of the polymerase activity of Cas10 (Santiago-Frangos et al., 2021), using two guide RNAs to initiate strand displacement amplification (CRISDA) (Zhou et al., 2018), and the electronic measurement of genomic sequence binding to graphene field-effect transistors (Hajian et al., 2019). Although still a new technology, there are already CRISPR-based tests approved by the United States Food and Drug Administration to target SARS-CoV-2 (Abudayyeh & Gootenberg, 2021).

CRISPR-dx assays can generally be designed to target any sequence, but the criteria for designing optimal guide RNAs depend on the specific method and Cas enzyme used. Cas enzymes differ in their requirements for adjacent sequence motifs, the length of guide RNA required, and how mismatches between the guide RNA and target affect cleavage efficiency. For example, Cas12 enzymes target both ssDNA and dsDNA, but for dsDNA targets, a protospacer adjacent motif (PAM) (e.g. TTTV) must be present in the target sequence near where the guide RNA binds (Kellner et al., 2019). This requirement can restrict which portions of the genome can be targeted, unless the PAM sequence is added to the 5’ end of one of the primers during a preamplification step (Li et al., 2019). The optimal length of the guide RNA can be different for each Cas ortholog: Cas13a uses a 28nt guide RNA, Cas13b uses a 30nt guide RNA, and Cas12a uses a 20nt guide RNA (Kellner et al., 2019). Finally, how much of an effect a mismatch between the guide RNA and the target has depends on both the location of the mismatch and the Cas enzyme used. For example, Cas13a is more sensitive to mismatches at the 3^rd^ position, whereas Cas12b is more sensitive to mismatches in the 10^th^ to 16^th^ positions (Li et al., 2019).

In addition to a specialized guide RNA, many methods require designing primers for preamplification of target DNA to increase sensitivity enough for the method to be useful for detecting pathogens in clinical settings. Amplification of target DNA is often done using an isothermal technique such as RPA or LAMP to minimize the need for expensive equipment and make it easier to conduct tests outside of laboratories (Abudayyeh & Gootenberg, 2021). Various non-target sequences may have to be added to the primer sequences, such as the previously mentioned PAM sequence required by Cas12 when targeting dsDNA (Li et al., 2019). Cas13 orthologs detect and cleave RNA, so to detect a DNA target a T7 promoter must be added to the 5’ end of one of the primers used for preamplification to allow for transcription (Kellner et al., 2019). These modifications can affect how primers and loci are chosen when designing a CRISPR-dx diagnostic assay.

Development of CRISPR-dx assays typically require the design of a diagnostic CRISPR guide RNA (crRNA) that discriminates between target and non-target sequences and sequence-specific primers for amplification of the target region. The abundance of public WGS data, combined with the affordability of generating new sequences for novel variants, has the potential to make designing new diagnostic assays fast and reliable. However, finding optimal candidate regions can be difficult when the assay is applied to organisms with large genomes or populations with high sequence diversity. We developed the python package `krisp` to identify candidate regions and primers for development of CRISPR-dx assays as well as any other RNA or DNA based assay. `Krisp` takes as input either a set of unaligned FASTA files representing assembled genomes or a Variant Call Format (VCF) file containing variants called against a reference genome. For FASTA input, `krisp` breaks sequences into k-mers and applies sequential filtering steps to find diagnostic regions that distinguish a target group from all outgroups and are flanked by conserved regions where primers can be designed. For VCF input, `krisp` identifies clusters of diagnostic variants flanked by regions without variants and infers the sequence for each group by applying variants to the reference sequence. For both input types, primers can be designed automatically using Primer3 (Untergasser et al., 2012) and the candidate regions can be filtered based on the presence and quality of possible primers. Candidate regions are reported as either human readable alignments or tabular data, allowing the user to apply further processing steps in a custom pipeline. Finally, `krisp` is highly flexible and can be configured to search for regions and primers compatible with any DNA/RNA probe/primer diagnostics as well as specific CRISPR-dx assays, such as SHERLOCK or DETECTR.

### Design and Implementation

The `krisp` python package has two principal functions: `krisp_fasta` and `krisp_vcf`. `Krisp_fasta` is used to infer diagnostic sites from whole genome assemblies based on shared unique k-mers. `Krisp_vcf` infers diagnostic sites from VCF files by analyzing a sliding window of variants and doing localized sequence inference. Each of these functions have a command line interface that is installed along with the package. `Krisp` is open source and available on GitHub with a user guide and test data (https://github.com/grunwaldlab/krisp) and on the python PyPI package repository (https://pypi.org/project/krisp/).

### Krisp_fasta

The `krisp_fasta` command is designed to find all diagnostic regions differentiating one set of sequences from another. Sequences for both target and nontarget organisms are broken down into k-mers representing potential diagnostic regions and primer sites. These k-mers are filtered on the following criteria: 1) The presence of polymorphisms that distinguish the target group from the non-target group and 2) sequence conserved in the target group on either side of the polymorphic region where primers can be designed. `Krisp_fasta` accepts a FASTA file representing each sample. FASTA files can be passed to the command as an “ingroup” file or “outgroup” file, corresponding to target and non-target organisms. Results in comma-separated value (CSV) format are streamed to standard output or saved to a file. Optionally, a more human-readable alignment format can be saved to a text file as well.

The algorithm relies on extracting, sorting, and intersecting k-mers to find diagnostic regions. For each input FASTA file, all k-mers of length ‘A’ (short for amplicon) are extracted. Included in length ‘A’ are regions of length ‘F’ and ‘R’, corresponding to conserved regions where primers could be designed on the ends of the amplicon. ‘F’ and ‘R’ surround a diagnostic region of length ‘D’, such that A = F + D + R (Figure 1). K-mers of length ‘A’ are extracted and sorted by the sequence content in the ‘F’ and ‘R’ regions. Sorted k-mers for each input FASTA file are written to intermediate files in parallel, along with the name of the file they came from. Pairs of k-mer files are then read in tandem and combined into new files of sorted k-mers that include the names of the files each k-mer was observed in. Pairs of k-mer files are processed in parallel. This process is repeated with its own output, combining pairs of files, until a single file is left that includes the sorted k-mers for all input FASTA files. In this file, k-mers that share identical ‘F’ and ‘R’ regions, but differ in the ‘D’ region, are grouped together since they were sorted previously. Each of these groups of consecutive k-mers is referred to as an alignment. Alignments with a single k-mer present in all the target group files and not present in any of the non-target group files are considered diagnostic. Primer3 is then used to search for potential primer sites in these diagnostic k-mers. Alignments with diagnostic k-mers and predicted primers are formatted for output in two formats: a human-readable alignment format in a plain text file and a CSV file for further bioinformatic analysis. Both output formats contain the sequence of the diagnostic region and the output of Primer3 for the best primers found.

**Figure 1.**
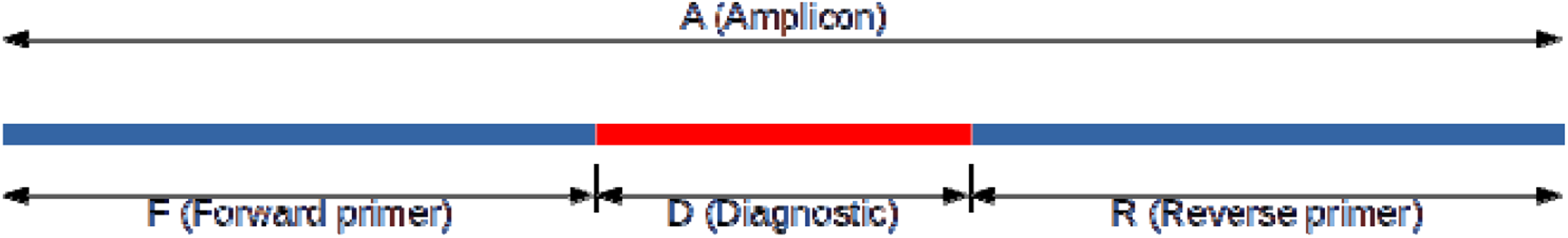
Definition of amplicon, diagnostic, and primer regions for developing candidate CRISPR-Cas assays with `krisp_fasta`. The total amplicon length A is subdivided into non-overlapping regions of length F, D, and R, such that A = F + D + R, where F and R correspond to the forward and reverse primer regions, and D corresponds to the diagnostic region.

### Krisp_vcf

The `krisp_vcf` function looks for clusters of diagnostic variants, referred to here as diagnostic regions, flanked by regions without variants where primers can be designed. Diagnostic variants are those that are conserved and exclusive to the target group. Variants are read from a VCF file with an associated reference file in FASTA format. A CSV file must be supplied that encodes which samples in the VCF belong to which group. Diagnostic regions can be discovered for two or more groups with a single execution of `krisp_vcf`. VCF data can be streamed from standard input or read from a file. The output of `krisp_vcf` can be streamed to standard out or written to a file. Additionally, a log containing progress and error messages can be streamed to standard error or written to a file.

`Krisp_vcf` uses a sliding window analysis of consecutive variants for each group of diagnostic regions evaluated. First, each chromosome (i.e., each sequence in the reference FASTA) is broken up into chunks to enable parallel processing, unless VCF data is being streamed from standard input, in which case parallel processing is not possible. For each chunk, variants are read in order and filtered by number of reads, number of samples, genotype quality, and other quality control metrics. For each group, variants passing these filters are supplied to a series of first in, first out (FIFO) queues representing the upstream region where the reverse primer will be designed, the diagnostic region where the crRNA will be designed, and the downstream region where the forward primer will be designed. When a variant is added to any of these queues, the sequence length spanned by the variants is inferred and if this is longer than a set value, variants from the queue are moved to the next queue until the inferred sequence length is short enough. The diagnostic region queue holds the maximum number of variants that can fit in the length of a crRNA and the maximum length of the primer queues is determined by the maximum amplicon length. Once a variant is removed from all of the queues, it is removed from memory, so only variants present in the queues contribute to the RAM usage of the program. Once all the queues have been filled, each time a variant is added to the series of queues described above, the variants in the queues are subjected to a series of tests to determine if they represent a diagnostic region. This effectively scans all possible regions in all samples. The following conditions must be met in order for the region to be considered diagnostic: 1. The diagnostic region queue must have enough diagnostic variants, 2. the variants in the diagnostic region queue must be conserved in the group of interest, 3. there must be enough conserved sequence in the group of interest in the primer queues to design primers, 4. A suitable primer pair must be found by Primer3 within the sequence inferred from the primer queues. These checks are ordered such that the most commonly failed and fastest to execute are done first in order to minimize processing time.

For regions passing all checks, the consensus sequence for the group of interest is inferred by applying the variants to the reference sequence and returned in the output. The output takes the form of a CSV file with columns for which group the region/primers is diagnostic for, the chromosome it occurs on, the coordinates of the reference genome where the primers and diagnostic region occur, the inferred sequence of the amplicon and surrounding sequence, and the Primer3 output. This format allows for additional downstream analysis with programming languages or spreadsheet programs, including giving users the sequence needed to design their own primers manually if desired. In addition to the CSV output, a human-readable text-based alignment output is provided that contains a multiple sequence alignment of the consensus sequence for each group being distinguished with annotations for the primer and crRNA locations. All Primer3 output is included in both the CSV and alignment-based formats.

### Performance evaluation

The performance of `krisp_fasta` was tested using a dataset of 12 assembled yeast genomes downloaded from NCBI: 6 genomes of baker’s yeast (*Saccharomyces cerevisiae*), and 6 genomes of the closely related budding yeast (*Saccharomyces kudriavzevii*), each roughly 10-12 mega-bases in length (Table 1). The effect of the number of cores used on run time was tested on two computing systems, a desktop with a 3.20GHz 6-core Intel i7-8700 CPU and 32GB of ram, and a computing cluster with a 2GHz 64-core AMD EPYC 7992 CPU and 4GB of ram. For both systems, `krisp_fasta` was instructed to find all genomic regions which distinguish these two species with a diagnostic region of length 10 and conserved primer region of length 20. The effects of the number of samples, mean genome length, and the length of the amplicon (i.e., k-mer) on run time and RAM were also evaluated on a laptop computer with an Intel® Core™ i7-10875H CPU @ 2.30GHz × 16 processor.

**Table 1.**
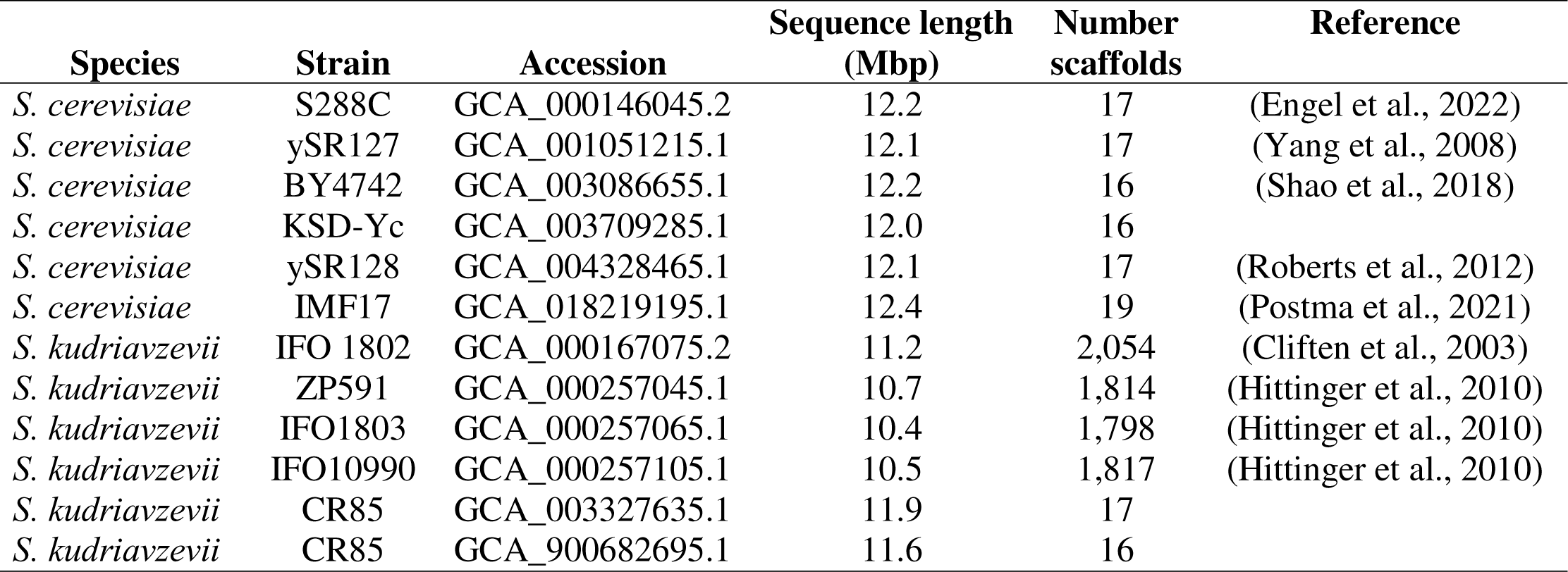
*Saccharomyces* genomes used to validate the krisp algorithm.

The performance of `krisp_vcf` was evaluated using a 20Gb VCF file containing 174,743,308 variants for 656 *P. ramorum* samples, grouped by clonal lineage. The 57.5 Mb reference genome of strain PR-102_v3.1 was used (Carleson et al., 2022). These tests were conducted on a laptop computer with an Intel® Core™ i7-10875H CPU @ 2.30GHz × 16 processor. The effect of the number of cores on execution time was evaluated on a subset of this dataset. The effects of number of samples, the number of groups compared, and the number of variants were evaluated using the same dataset and computer by subsetting the dataset as needed.

### Validating krisp_fasta and krisp_vcf

The output of `krisp_fasta` and `krisp_vcf` were validated in the lab by using both to design a SHERLOCK diagnostic assay to differentiate the plant pathogen *Phytophthora ramorum* from closely related *Phytophthora* species (Grünwald et al., 2019). For `krisp_fasta` whole genome sequences from 5 *P. ramorum* isolates (PR-102, PR-15-019, PR-18-069, PR-18-108, and PR-18-126) were used to represent the diversity within *P. ramorum* and genome sequences of *P. brassicae*, *P. cryptogea*, *P. foliorum*, *P. hibernalis*, *P. lateralis*, and *P. syringae* were used to represent non-target *Phytophthora* species. In addition, a version of the *P. ramorum* genome PR-102 with variable regions masked with Ns was included. The masking was done by analyzing a VCF file containing published variants of *P. ramorum* samples (Dale et al., 2019; Kasuga et al., 2016; Sambles et al., 2015) and converting any position homozygous for the alternative allele to N. This allowed for the incorporation of data from many samples that do not have assembled genomes available. For `krisp_vcf`, a VCF file containing variants of *P. ramorum* samples from available genomes (Dale et al., 2019; Kasuga et al., 2016; Sambles et al., 2015) and the PR-102_v3.1 reference sequence was used as input (Carleson et al., 2022). A promising diagnostic site was selected from the many options produced by `krisp_fasta` and checked against analogous VCF data using `krisp_vcf`.

Briefly, we evaluated the ability to distinguish *P. ramorum* from other *Phytophthora* taxa with a SHERLOCK assay. SHERLOCK is a two-step assay starting with RPA of target DNA. The forward Primer for RPA included a T7 promoter region at the 5’ end that, when combined with the reverse primer, produced a 117 bp amplicon in the target region (Forward primer: GAAATTAATACGACTCACTATAGGGTGCATTTTCGACAAATTCGAGTGCGGGGTCAG, Reverse primer: ATCGAAATATCGGCGCGTCCATAACGGTCATA). Amplification reactions were prepared according to Kellner et al. (2019), using a master mix that included 10µM primers, water, and TwistAmp rehydration buffer (Twistdx, Maidenhead, UK). This master mix was used to rehydrate the TwistAmp polymerase followed by the addition of 280mM of magnesium acetate. Ten microliter aliquots of the reconstituted RPA were added to PCR tubes along with 1µl of template and placed in a thermocycler set at 37C for 30 min. The crRNA 5’-UUAUCCGAGCCCGUGAUGAAGUUGUUGC-3’ was designed for the Protein phosphatase 2 (PP2A) regulatory subunit B locus. The 5th base was modified from an ‘A’ to a ’C’ to introduce 1 mismatch into the crRNA-target alignment. For *P. ramorum*, this meant that the crRNA and target have a single mismatch. For other *Phytophthora* species, there were at least two mismatches and thus no collateral cleavage should occur. Adding the conserved DR region for LwaCas13a results in 5’-ACUACCCCAAAAACGAAGGGGACUAAAACUUAUCCGAGCCCGUGAUGAAGUUGUUGC-3’. For detection, a master mix was prepared containing ultrapure water, 20mM HEPES pH 6.8, 9.5mM MgCl, 1mM rNTP solution mix, 6.7µg LwaCas13a (MCLAB, San Francisco, CA or Genscript, Piscataway, NJ), 40U Murine RNase inhibitor (New England Biolabs, Ipswitch, MA), 2.5U T7 RNA polymerase (New England Biolabs), 10ng/µl CRISPR guide RNA (IDT, Coralville, IA), and 0.13µM RNaseAlert v2 (Thermo Fisher, Waltham, MA) per reaction. Four replicates of each sample were aliquoted into a 384 well plate (20µl per well), centrifuged and immediately placed in a fluorescent plate reader (Tecan, Switzerland) preheated to 37C. Fluorescence (490/520nm) was recorded for 3h at 5min intervals. Positive controls included known samples of *P. ramorum* from clonal lineages NA1, NA2, EU1 and EU2, whereas negative controls included samples of *Phytophthora* species including *P. cinnamomi, P. foliorum, P. lateralis,* and *P. plurivora* as well as water. Background subtracted fluorescence was graphed over time.

## Results

`Krisp` is a python package for finding candidate regions for the development of CRISPR-dx diagnostic assays. `Krisp` can analyze whole genome sequence data in the form of FASTA files or variant data in the form of a VCF file with an associated reference in FASTA format. Primer3 is used to screen potential diagnostic regions for suitable primer binding sites. Diagnostic regions are output in the form of a CSV file or human-readable plain text alignments. `Krisp` has been optimized to minimize RAM use and can run in parallel, allowing large datasets to be processed on personal computer in a matter of hours.

### Expected performance

We estimated the theoretical expected performance of `krisp_fasta`. The first computationally intensive step is the extraction of candidate regions as k-mers from genome files and sorting by the conserved primer regions. The extraction of k-mers takes *O(S*A*N)* time, where *S* is the number of samples, *A* is the amplicon length, and *N* is the average number of bases in a sample’s sequence. The subsequent sorting step takes *O(S*A*N*log(N))* time because each genome file is expected to contain ‘N’ potential amplicons, which require *O(A*N*log(N))* comparison and sorting operations. Since sorting is computationally slower, the extraction and sorting of k-mers has an expected runtime complexity of *O(S*A*N*log(N)*). This means that the compute time is expected to scale linearly with the number of genome files and the amplicon length, and log-linear with mean sequence size. The other computationally intensive step is calculating the intersection of the k-mers between groups of samples. Since these k-mers are already sorted, finding conserved primer pairs can be done with a single traversal through the k-mer files. The time complexity of this step is roughly *O(S*A*N)*, as we iterate through roughly *N* amplicons and compare *S* potential amplicons across *A* different sites. This means that compute time for this step is expected to scale linearly with the number of samples, amplicon length, and mean sequence length.

We also estimated the theoretical expected performance of `krisp_vcf`. Since `krisp_vcf` reads all variants only once, processing time scales linearly with number of variants. It also should scale linearly with the number of groups being distinguished (G) since each group gets its own set of processing queues and quality filters. Finally, it should also scale linearly with the number of samples in the VCF file (S). This results in a predicted time complexity of O(V*G*S).

When multiple cores are available for computation, the real-world runtime can be significantly reduced since many of these steps are independent, or nearly independent. For example, extracting and sorting k-mers from the input files are completely independent tasks and are executed as such by `krisp_fasta`. Similarly, taking the intersection of k-mers is done in parallel by repeatedly taking the intersection of pairs of sample files, thereby building an inverted binary tree where the root node (i.e., the final file produced) is the intersection of all samples. Each intersection job is executed in parallel as a separate task, and each task is further written in parallel to utilize multiple cores. `Krisp_vcf` also takes advantage of parallel processing by splitting the input VCF file into chunks based on reference sequence coordinates, allowing for each chunk to be processed in parallel.

### Observed performance

On both computing systems, `krisp_fasta` ran about twice as fast when 6 cores were used. Further increasing the number of cores on the desktop computer provided no increase in speed, likely due to hardware limits being reached. When run on the computing cluster, speed increased through 12 cores (Figure 2). In terms of absolute time, the desktop computer took ∼40 minutes to complete processing the test dataset with 1 core, which decreased to ∼20 minutes when using 6 cores. The computing cluster utilizes a slower CPU and took ∼80 minutes to complete with 1 core, which decreased to ∼25 minutes with 12 cores (Figure 2). The `krisp_fasta` algorithm utilizes very little memory during execution, since it uses intermediate files instead of RAM for memory-intensive steps. Although RAM usage appears to correlate with input sequence length (Supplementary Figure 1), this is due to the Linux `sort` utility taking advantage of excess RAM to operate faster. If less RAM were available the program would still run. `Krisp_vcf` achieves a low memory footprint as well by only loading variants present in a sliding window. This allows `krisp` to run on computing systems with as little as 4GB of RAM, making it suitable for laptops, desktops, and similar personal computers. The effects of the number of samples, mean genome length, and the length of the amplicon (i.e. k-mer) on run time and RAM were confirmed to be linear as expected (Supplementary Figure 2).

**Figure 2.**
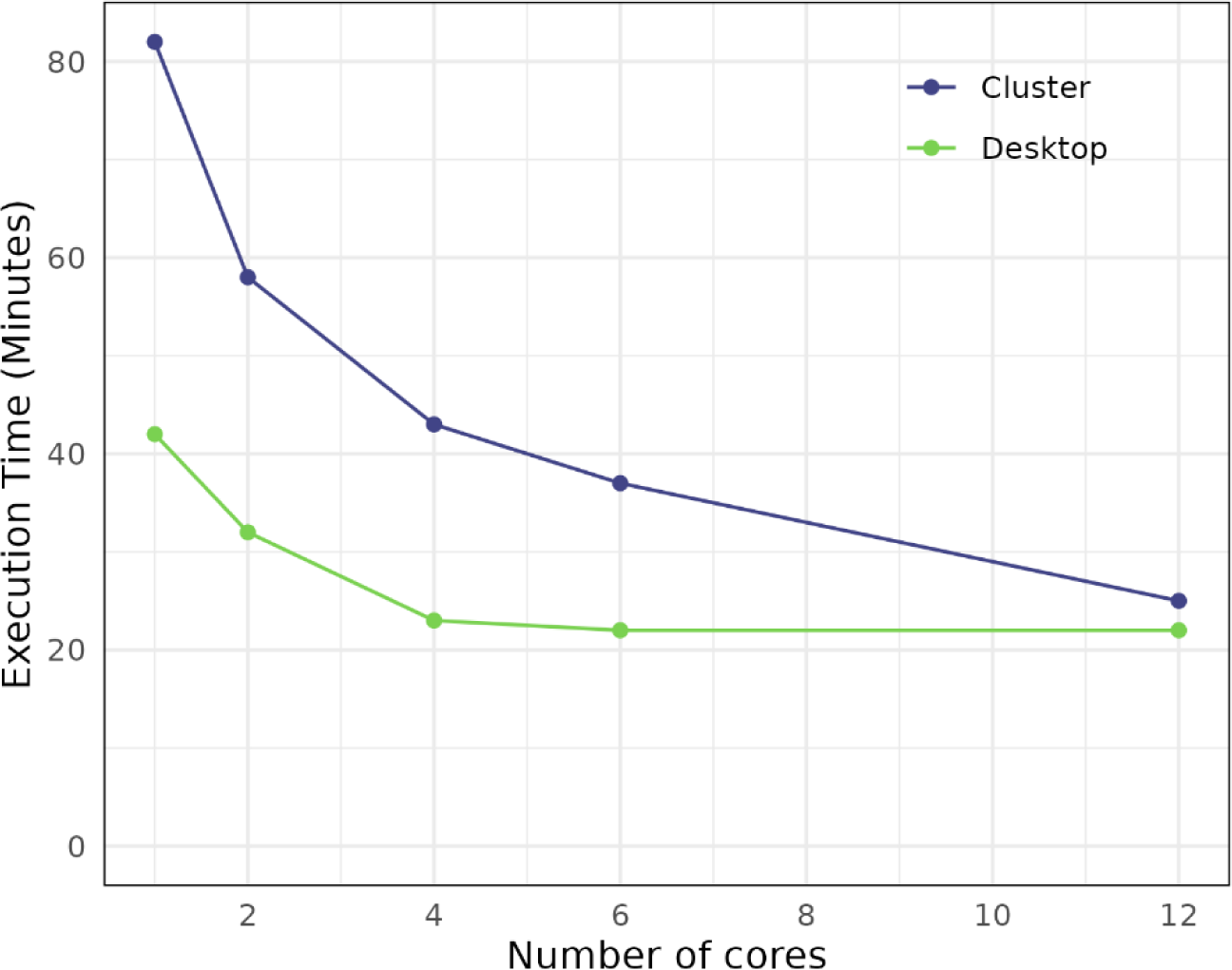
Execution time of `krisp_fasta` on a test dataset of 12 yeast genomes with varying numbers of processors used. See Table 1 for more information on the data used. Desktop timing tests were performed on a 6-core Intel i7-8700 CPU with a clock rate of 3.20GHz and 32GB of ram. Cluster timing tests were performed on a 64-core AMD EPYC 7992 CPU with a clock rate of 2GHz and 4GB of ram.

`Krisp_vcf` completed analysis of an entire 20Gb VCF file containing 13 million variants for 656 samples in 24 minutes when using 16 threads on a laptop. The effect of the number of threads used on the execution time was tested using a subset of the same data. Increasing the number of cores from 1 to 4 decreased execution time nearly 4-fold, but additional cores had little effect, likely due to limitations of the laptop cooling system (Figure 3). The effects of number of samples, the number of groups compared, and the number of variants on execution time was confirmed to be approximately linear, although the trend was somewhat noisy (Supplemental Figure 2). RAM usage was consistently around 170Mb regardless of dataset scale.

**Figure 3.**
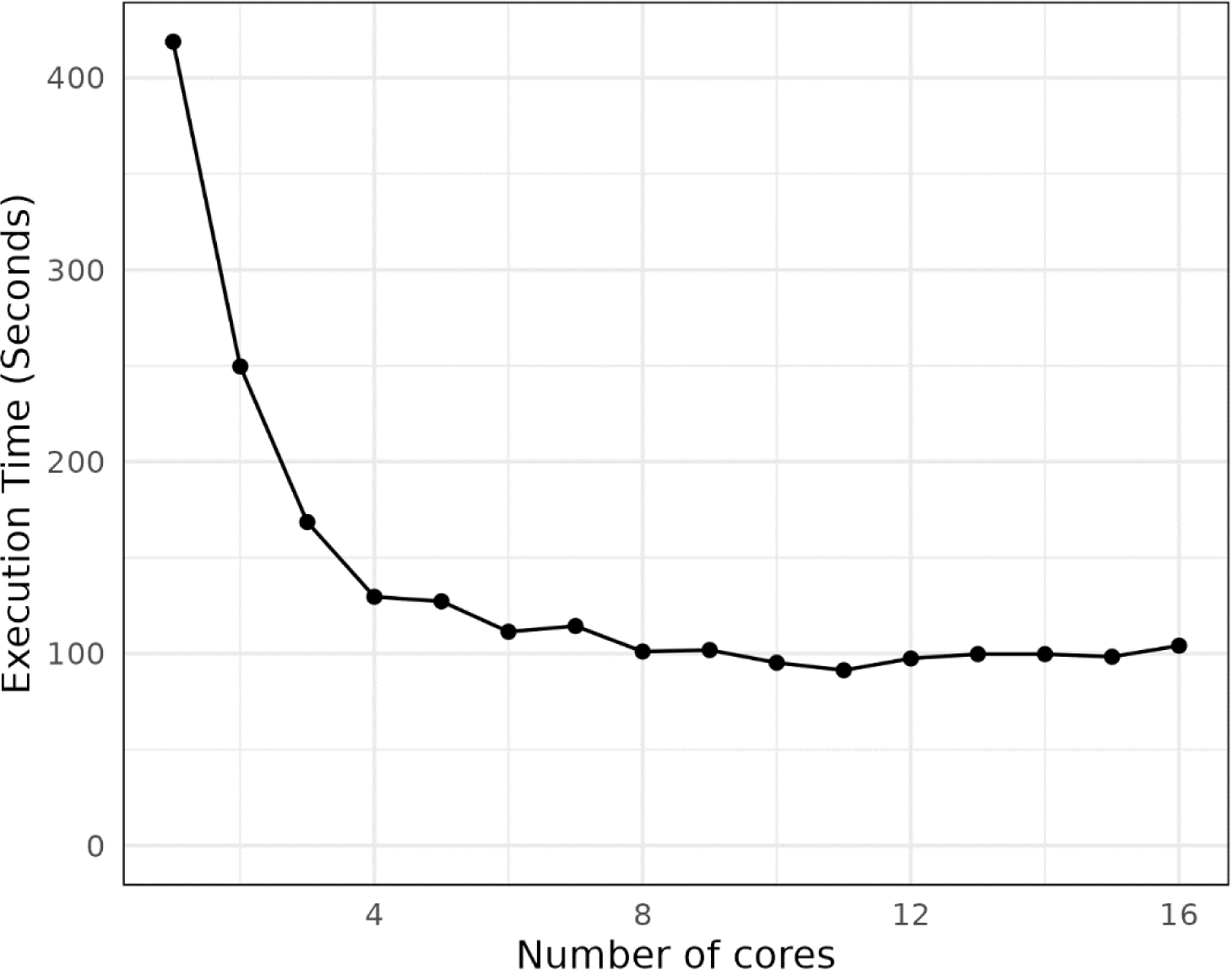
The effect of the number of cores used on the execution time of `krisp_vcf`. The test dataset used consisted of 10 samples with 522,965 variants distinguishing 2 groups. These tests were done on a laptop computer with an Intel® Core™ i7-10875H CPU @ 2.30GHz × 16 processor.

### Laboratory validation

The outputs of both `krisp_vcf` and `krisp_fasta` were used to design a SHERLOCK proof-of-concept assay (Figure 4, top). The assay was built to distinguish *Phytophthora ramorum* from other closely related *Phytophthora* species. Our results suggest that both programs work as expected. All *P. ramorum* variants NA1, NA2, EU1 and EU2 resulted in fluorescence whereas negative controls, including 6 other *Phytophthora* species, did not (Figure 4).

**Figure 4.**
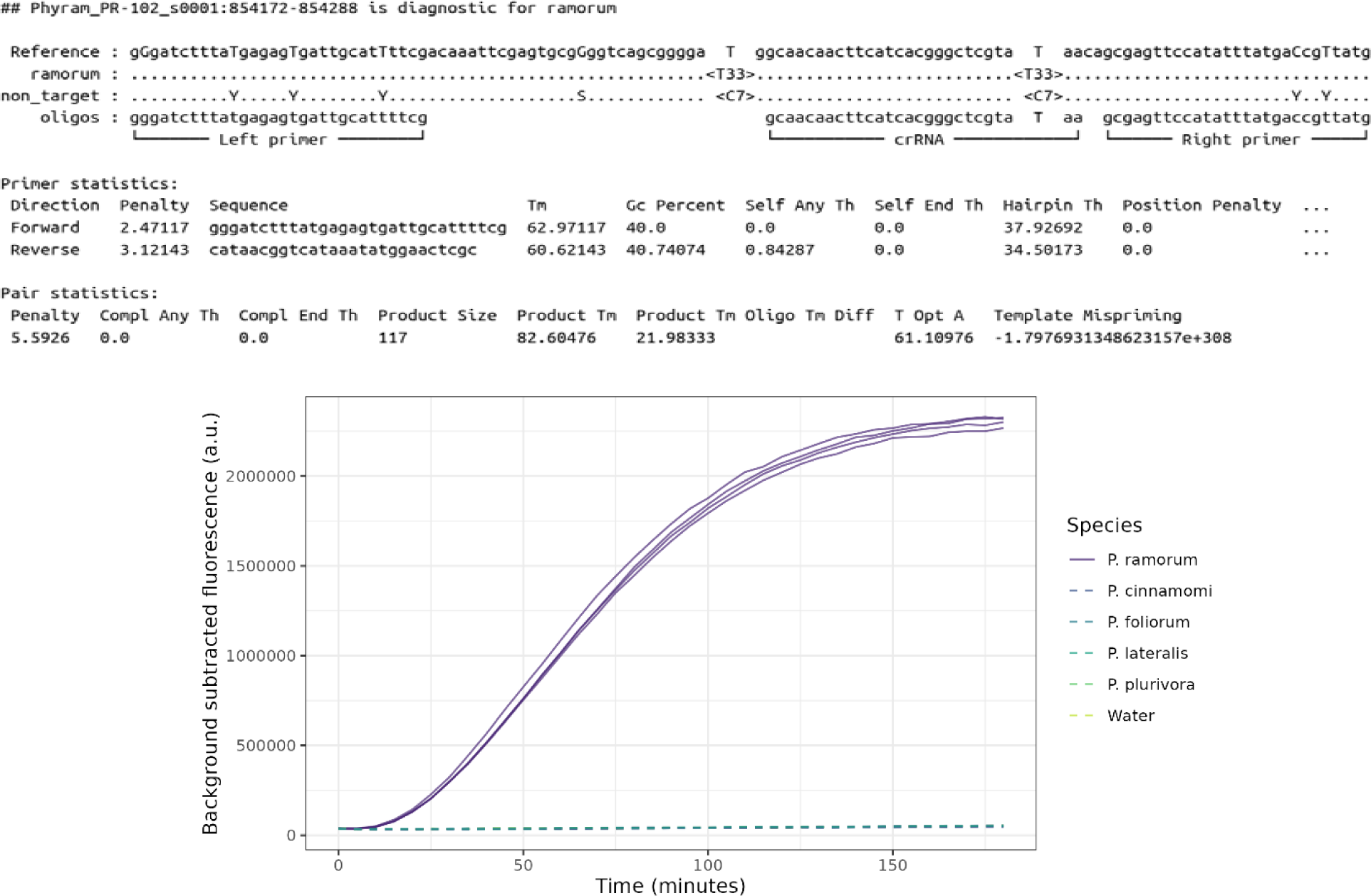
Proof-of-concept predicted by both `krisp_vcf` and `krisp_fasta` to create a SHERLOCK CRISPR-dx assay for the Protein phosphatase 2 (PP2A) regulatory subunit B locus to detect the species *Phytophthora ramorum.* **Top graph:** Abbreviated output from `krisp_vcf` showing predicted primers and diagnostic crRNA region for detection of *P. ramorum*. Note, that the primers we initially used for validation were different, but the crRNA locus is identical. **Bottom graph:** *P. ramorum* clonal lineages NA1, NA2, EU1, EU2 as well as Asian strains (purple solid lines) could be detected while non-target species *Phytophthora cinnamomi, P. foliorum, P. lateralis,* and *P. plurivora* (shown in various colors as dashed lines) as well as other groups including fungi and bacteria (not shown) did not amplify.

## Discussion

The ability to rapidly develop diagnostic tests to distinguish closely related organisms would be a useful tool for tracking the spread of emerging pathogens and organizing an effective response. CRISPR-dx technology has the potential to produce highly specific, sensitive, and inexpensive tests that can be administered with minimal specialized equipment or expertise. Quickly identifying candidate sequences to design diagnostic crRNAs and primers that can be tested in the laboratory will decrease the time it takes to deploy new assays. `Krisp` can be used to find candidate diagnostic regions in which to design CRISPR-dx or amplification-based assays to differentiate one group of organisms from another for any species for which assembled sequences or variant data is available. `Krisp` has been carefully optimized to handle large datasets of commonly available data formats and leverage parallel computing to rapidly produce actionable results.

To demonstrate the usage of `krisp_fasta`, below we show how it can be used on the yeast dataset described above to locate regions where a CRISPR-dx assay could be designed (Table 1). We aim to find diagnostic regions which differentiate the 6 baker’s yeast genomes from the 6 budding yeast genomes. More specifically, we instruct `krisp` to find all genomic regions in which a diagnostic region of length 10 is flanked by conserved primer regions of 50 nucleotides in both directions:

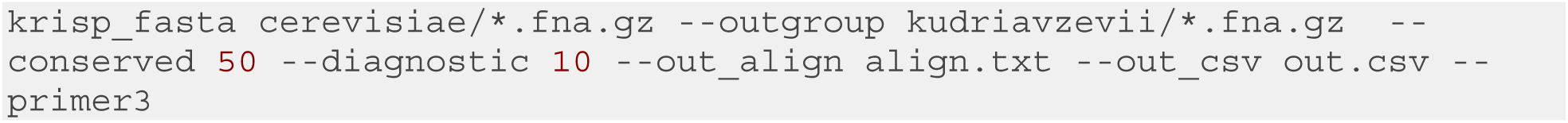

In this example, `krisp_fasta` was able to find 120 candidate regions which fit the criteria. Below is one of those regions (parts of the output are abbreviated with ‘…’ for visualization):

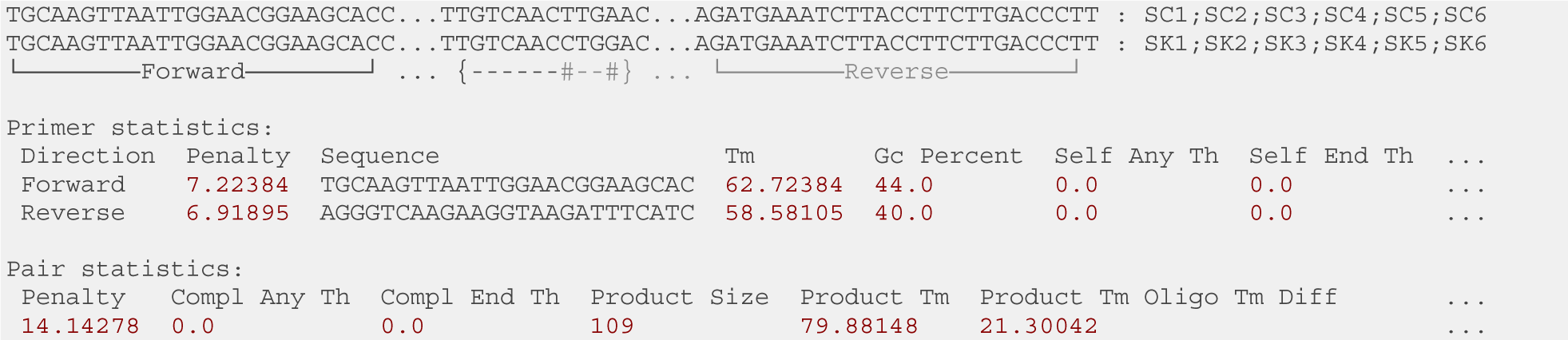

Candidate regions can be output in an alignment format where each unique sequence is stacked vertically and the corresponding genome files are listed on the right. In this example, there are two sequences and twelve genome files with names starting with ‘SC’ or ‘SK’, corresponding to the genome files for *S. cerevisiae* and *S. kudriavzevii*, respectively. The first sequence is associated with six genome files, all of which correspond to *S. cerevisiae*, implying that this sequence is conserved across the entire ingroup, whereas the second sequence was found exclusively in the outgroup *S. kudriavzevii*. Since only unique sequences are displayed, it is common to have multiple files associated with a single sequence in an alignment. When a sequence is found multiple times in a genome file, the number of times it occurred is appended to the genome file name in the format of ‘(n)’. For example, ‘SC1(2)’ would imply that this sequence was detected twice in the genome file SC1. Near-identical sequence matches would be listed as separate sequence entries, assuming the only differences are within the diagnostic region. The last line of the alignment shows a summary, where ‘{}’ denotes the boundaries of the diagnostic region, ‘-’ a conserved position, ‘*’ a non-diagnostic SNP, and ‘#’ a diagnostic SNP with respect to the ingroup. In this case we see that the diagnostic region contains two diagnostic SNP’s, a ‘T-C’ and ‘A-G’ difference, and zero non-diagnostic SNP’s. The alignment is annotated with the locations of the best primers found by Primer3. The full output of Primer3 for these primers is printed in a tabular format below the alignment for convenient manual inspection.

In the case of VCF input, `krisp_vcf` can be used in a similar way to `krisp_fasta`. For this example, we will use a subset of the of *P. ramorum* VCF data described above that is included in the package as a test dataset. We search for regions that distinguish each of three clonal lineages of *P. ramorum* from all other lineages with the following command:

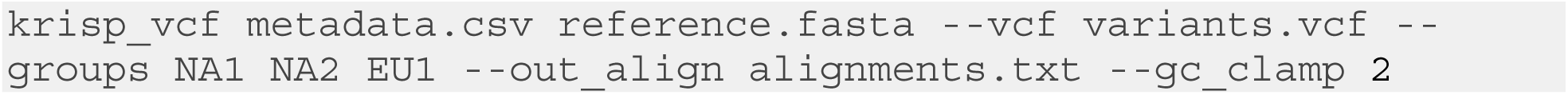

The “metadata.csv” file contains a table with two columns: each sample’s name as it appears in the VCF file and what group (lineage in this case) that the sample belongs to. The “--groups” option defines which groups assays should be designed for. The “--gc_clamp 2” option is one of many Primer3 settings that can be changed; this option instructs Primer3 to only consider primers with at least 2 bases that are G or C on the 3’ end of both primers. The command above produces the following alignment, among many others (parts of the output are shortened with ‘…’ for visualization):

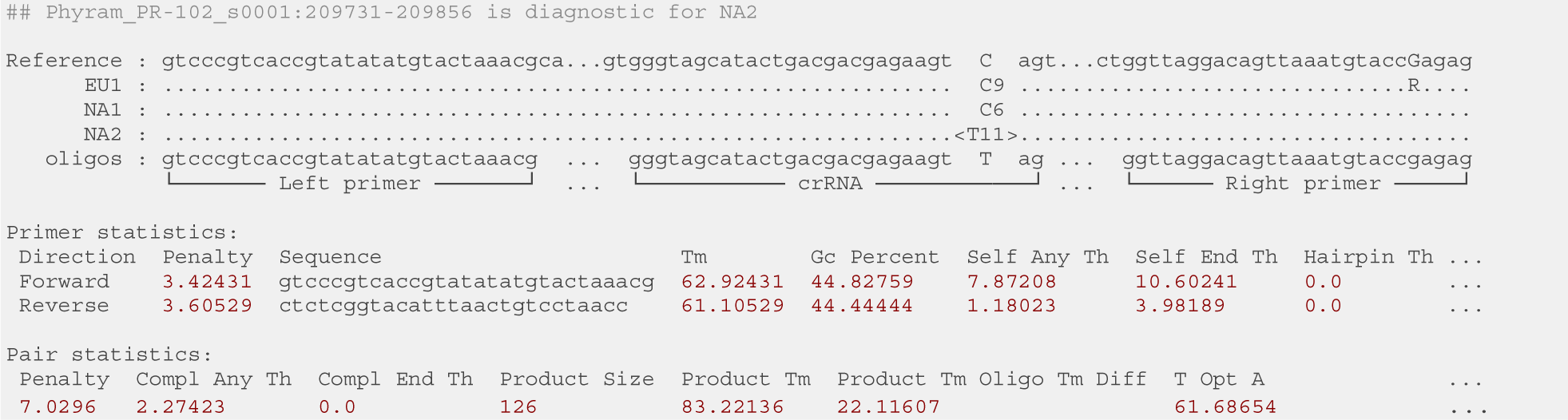

In this format, columns in the alignment with a diagnostic variant have the alleles and their counts listed in the form of the allele sequence followed by the number of samples that have that allele. For example, the “C9” in the above output means that all 9 of the samples assigned to the EU1 lineage have C in that position. If multiple alleles were present, they would be listed in series (e.g., C5T1). Diagnostic alleles are highlighted with angle brackets. For example, in the above output “<T11>” means that all 11 samples of NA2 have T at that position and none of the samples from other lineages do. Variants not located in diagnostic columns are indicated by capital letters, potentially using IUPAC ambiguity codes if a group has multiple alleles at a given position. Like `krisp_fasta`, the Primer3 output for the best primers is presented below the alignment.

We hope that the `krisp` algorithm will facilitate development of novel diagnostic assays. A complete step-by-step design of a CRISPR-dx assay is beyond the scope of this paper; here, we provide a starting point for more detailed and customized analyses. The designing of a typical CRISPR-dx assay can be broken down into two steps, corresponding to primer-based amplification and CRISPR-Cas based recognition via a crRNA. `Krisp` is designed to provide possible candidates for both, but final selection of a crRNA and primers will depend on many factors, including the Cas enzyme and type of amplification (e.g., RPA or LAMP) used. For example, the LwCas13a enzyme used in SHERLOCK can only achieve single base pair resolution when the SNP is in the 3^rd^ position from the 3’ end and an artificial mismatch is added nearby. `Krisp_vcf` takes this into account by positioning the diagnostic SNPs at a location specified by the user. We hope to add additional functionality to `krisp` in the future to consider such technique-specific details to further decrease the workload needed to design CRISPR-dx or other diagnostic assays. Finally, `krisp` can be used beyond CRIPSR-dx applications to search genome or VCF data for any combination of user specified diagnostic probes and/or primers distinguishing target and non-target groups.

## Availability and future directions

Krisp is open source and available on GitHub (https://github.com/grunwaldlab/krisp) and the python PyPI package repository (https://pypi.org/project/krisp/). It is released under the MIT license. A user guide with examples is provided on the GitHub repository.

## Acknowledgements

This work was supported by grants from USDA ARS (2072-22000-041-000-D), USDA NIFA (2023-67013-39918) and USDA APHIS to NG.

---

**Supplementary Figure 1.**
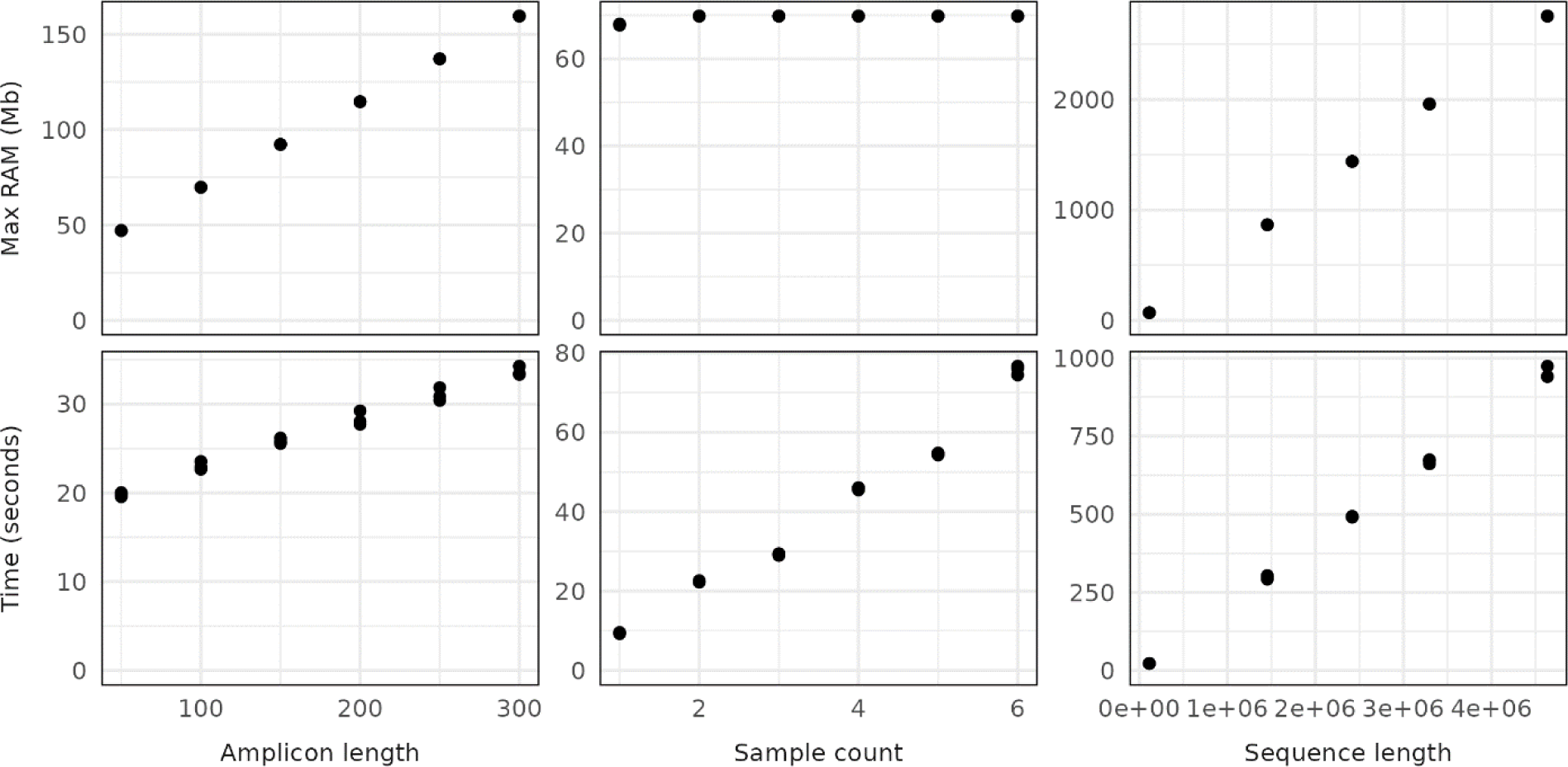
The effects of sample count, amplicon (k-mer) length, and mean genome length on the execution time and maximum RAM usage of `krisp_fasta`. For each column, variables not being tested were held constant with the following values: sample count of 2, amplicon length of 100bp, and a mean sequence length of 69,808 (the length of the first chromosome in the test dataset). Testing was done a laptop computer with an Intel® Core™ i7-10875H CPU @ 2.30GHz × 16 using a single core.

**Supplementary Figure 2.**
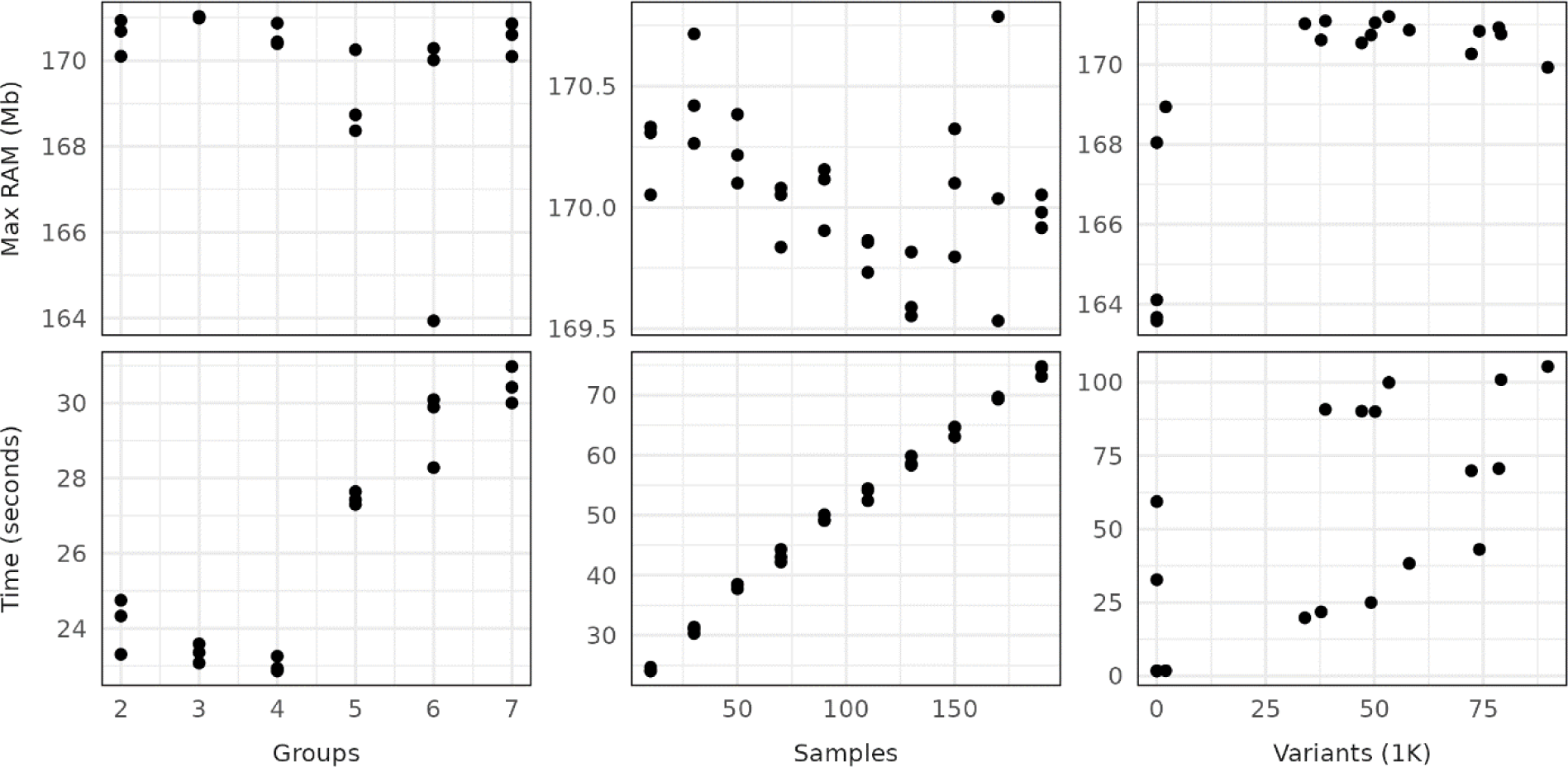
The effects of variant count, sample count, and number of groups being distinguished on the execution time and maximum RAM usage of `krisp_vcf`. For the sample count and group count tests, variant count was held constant at the number of variants that occur in the first 500,000bp of the first chromosome. For the variant count and group count tests, sample count was held constant at a total of 6 and 24 samples respectively. For the variant count and sample count tests, the number of groups was held constant at 2. Testing was done a laptop computer with an Intel® Core™ i7-10875H CPU @ 2.30GHz × 16 using a single core.

